# Host transcriptome and microbiome interactions in Holstein cattle under heat stress condition

**DOI:** 10.1101/2022.10.04.510823

**Authors:** Bartosz Czech, Yachun Wang, Kai Wang, Hanpeng Luo, Lirong Hu, Joanna Szyda

## Abstract

Climate changes affects animal physiology. In particular, rising ambient temperatures reduce animal vitality due to heat stress and this can be observed at various levels which included genome, transcriptome, and microbiome. In a previous study, microbiota highly associated with changes in cattle physiology were identified under heat stress conditions which included rectal temperature, drooling score, and respiratory score. In the present study, genes differentially expressed between individuals were selected representing different additive genetic effects towards the heat stress response in cattle in their production condition. Moreover, a correlation network analysis was performed to identify interactions between the transcriptome and microbiome for 71 Chinese Holstein cows sequenced for mRNA from blood samples and for 16S rRNA genes from fecal samples. Bioinformatics analysis was performed comprising: i) clustering and classification of 16S rRNA sequence reads, ii) mapping cows’ transcripts to the reference genome and their expression quantification, and iii) statistical analysis of both data types – including differential gene expression analysis and gene set enrichment analysis. A weighted co-expression network analysis was carried out to assess changes in the association between gene expression and microbiota abundance as well as to find hub genes/microbiota responsible for the regulation of gene expression under heat stress. Results showed 1,851 differentially expressed genes were found that were shared by three heat stress phenotypes. These genes were predominantly associated with the cytokine-cytokine receptor interaction pathway. The interaction analysis revealed three modules of genes and microbiota associated with rectal temperature with which two hubs of those modules were bacterial species, demonstrating the importance of the microbiome in the regulation of gene expression during heat stress. Genes and microbiota from the significant modules can be used as biomarkers of heat stress in cattle.

## 1 INTRODUCTION

Humans induce global warming that negatively influences all organisms on the Earth, e.g. by the occurrence of heat stress (HS) in livestock, which negatively affects its vitality, physiological responses, and behavior. Heat stress can inhibit milk production in dairy cattle which can result in significant losses to the industry (Garner et al., 2020). Moreover, HS in cows results in further economic losses by reducing reproduction (Macciotta et al., 2017). Currently, phenotypes such as rectal temperature, drooling score and respiration rate score are standard physiological indicators of HS (Luo et al., 2021; Brito et al., 2020). Currently, due to genomic selection being targeted to increased milkr production in cattle, cows tend to be more susceptible to HS (Biffani et al., 2016). The phenomenon of HS from the molecular perspective is a complex challenge and still, many mechanisms are unknown. A previous study focusing on the differential abundance of microbiota already demonstrated that HS affects the microbial composition of the colon (Czech et al., 2022). Only a few studies have demonstrated the effect of HS on the transcriptome profile of cattle, and have identified genes potentially associated with the HS response. Gao (Gao et al., 2019) pointed out that amino acid and glucose transport were downregulated by HS. In another study looking at the effect of HS on the transcriptome profile of mammary glands of cows (Yue et al., 2020), the authors indicated that HS affects dairy cows’ immunity and thus has a potential impact on milk yield. Sigdel *et al*. also presented the association analysis of HS cattle using SNP markers from the Cooperative Dairy DNA Repository and the Council on Dairy Cattle Breeding, and identified genes *HSF1, MAPK8IP1*, and *CDKN1B* that were directly involved in the cellular response to HS (Sigdel et al., 2019).

In general, the impact of HS on cows is fairly difficult to assess due to the complexity of the metabolism and physiology of cows. However, the development of molecular techniques like next-generation sequencing, mass spectrometry, and other techniques allow us to look more deeply into these mechanisms by obtaining information about the entire biology of the system. Additionally, high-performance computers with new algorithms allow for considering more complex statistical models that allow for better insights into the complexity of organisms (Park et al., 2021b). Many other studies that focused on the integration analysis of host transcriptome and microbiota already demonstrated the importance of the multiomics approach to identify biomarkers underlying diseases and complex traits (Wang et al., 2019). In livestock, only a few studies have been focusing on the integration of host transcriptome and microbiome interactions. One of the studies showed the impact of the interaction of host transcriptome and microbiome on the physiology of full-sibs broilers with divergent feed conversion ratio (Shah et al., 2019). Ramayo-Caldas et al. (Ramayo-Caldas et al., 2021) investigated the joint effects of host genomic variation and the gut microbiome variation in the context of immune response in pigs. Also, Carillier-Jacquin et al. (Carillier-Jacquin et al., 2022) considered the importance of using both sources of information for the accuracy of prediction of pig digestibility coefficients, concluding that the incorporation of gut microbiome information is important for prediction and even outperform the importance of host genetic variation. Another study in chickens, showed the influence of nutrition on the interaction between transcriptome and microbiome which in turn influenced egg production in aged laying hens (Liu et al., 2022). Although those studies have revealed (and stressed) the importance of the incorporation of microbiome information into the evaluation of phenotypes that are important for livestock, the particular impact of the interaction of host transcriptome and microbiome on HS is still not well understood. Moreover, previous studies (Freitas et al., 2022) indicated a complex genetic architecture underlying the HS response, so it is expected that using all available sources of omic information is crucial for the modeling of this phenotype.

Therefore, it is worth mentioning that almost all HS studies used the case-control experimental design. In this study, we used a continuous variable to measure the HS response, to reflect the production environment which allowed us to study potential changes in gene expression level, microbiome abundance, and their interactions under standard conditions. This study aimed to identify genes differentially expressed between cows characterized by different additive genetic effects of HS response measured by the three HS indicators: rectal temperature, drooling score, and respiratory score, and to perform an integration of mulitomics data of host gene expression levels in relation to its microbiome composition.

## 2 MATERIAL AND METHODS

### 2.1 Material

Fecal and blood samples from 71 Chinese Holstein cows were collected in 2017, 2018, and 2019. Cows were sampled once over the course of three years. In this study no artifical heat stress challenge was imposed since the major goal was to assess the impact of heat stress that occurs during standard production conditions and is due to a combination of several climatic and production factors. Heat stress phenotypes used in this study were represented by the additive genetic effect of each cow, corrected for environmental factors such as lactation stage, age at calving, parity, and temperature-humidity index, that were expressed as deregressed estimated breeding value (DRP) for rectal temperature, respiratory score, and drooling score. We used a mixed linear model for the DRP estimation:

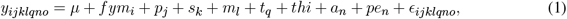

where *y*_*ijklqno*_ refers to phenotype (RS, DS or RT), *μ* is the population mean, *fym*_*i*_ is the fixed effect for farm-year, *p*_*j*_ is the fixed effect of parity, *s*_*k*_ is the fixed effect of lactation stage, *m*_*l*_ is the fixed effect of the indication if the animal is before or after milking, *t*_*q*_ is the fixed effect of testing time (morning or afternoon), *thi* is the fixed effect of temperature-humidity index, *a*_*n*_ is the animal additive genetic effect, *pe*_*n*_ is the permanent environmental effect, and *E*_*ijklqno*_ is the random residual. The covariance matrix of random effects has the following structure:

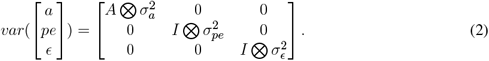

More information about the sampling procedure, housing of animals, phenotypes, THI, and statistical model used to estimate DRP and a description of the dataset were included in a previous study (Czech et al., 2022).

#### 2.1.1 Ethics approval and consent to participate

The data collection process was carried out in strict accordance with the protocol approved by the Animal Welfare Committee of the China Agricultural University. All experimental protocols were approved by the Animal Welfare Committee of the China Agricultural University. All methods are reported in accordance with ARRIVE guidelines (https://arriveguidelines.org) for the reporting of animal experiments.

### 2.2 Methods

#### 2.2.1 Microbiome

Deoxyribonucleic acid was isolated from fecal samples, which represent the microbiome composition of the colon and were used for sequencing of V3 and V4 regions of the 16S rRNA gene using the Illumina MiSeq and HiSeq platforms. Sequenced reads were cleaned and processed using QIIME2 software (Bolyen et al., 2019) with the SILVA database (Quast et al., 2012) to cluster and classify them to taxonomical levels. The final output is represented by the amplicon sequence variants table with the information about the frequency of a given taxon in a given fecal sample. The procedure is explained in detail by Czech et al. (Czech et al., 2022).

#### 2.2.2 mRNA-seq

Total RNA was isolated from from leukocytes according to the instructions of the TRIzol Reagent method (Rio et al., 2010). The cDNA library was prepared using mRNA molecules and sequenced using the NovaSeq 6000 System Illumina platform. Ribonucleic acid concentration and quality were determined using Equalbit RNA BR Assay Kit (Invitrogen, California, USA) and the Nanodrop 2000 (Thermo, Massachusetts, USA). Ribonucleic acid integrity was assessed using 1% agarose gel electrophoresis, and then used for library construction with 28S/18S *>*1. For the RNA-Seq library, 2 *μ*g total RNA was firstly used for purification and fragmentation with NEBNext Poly(A) mRNA Magnetic Isolation Module (Cat No. E7490S, New England Biolabs (UK) Ltd., Hitchin, Herts, UK) and then followed by cDNA library with NEBNext Ultra RNA Library Prep Kit for Illumina (Cat No. E7530S, New England Biolabs (UK) Ltd., Hitchin, Herts, UK). All libraries were quantitated by the Equalbit DNA BR Assay Kit (Invitrogen, California, USA) and pooled to generate equimolarly, and finally submitted for sequencing by the NovaSeq 6000 System (Illumina, Inc., San Diego, Californi, USA) which generated 150 base paired-end reads. Sequenced reads were evaluated in the context of their quality and cleaned using Fastp software (Chen et al., 2018). Filtered reads were mapped to the bovine genome (ARS-UCD1.2) using STAR software (Dobin et al., 2012) and Picard (Broad Institute, Accessed: 2022/07/15; version 2.27.4) was applied to mark duplicates. Finally, RNA-SeQC (DeLuca et al., 2012) software was used to quantify the expression. Gene expression was analyzed using the DESeq2 R package (Love et al., 2014) to perform differential gene expression analysis fitting the negative binomial regression model adjusted for the sequencing year. The effect of HS was expressed as the average fold change per DRP increased by one unit. The Wald test was used to assess the significance of slope estimates. P-values obtained separately for each gene were corrected for multiple testing using the Benjamini-Hochberg method (Benjamini and Hochberg, 1995) for controlling the False Discovery Rate (FDR). Genes with the FDR *<* 0.05 were considered to be associated with HS. Next, we performed Gene-Set Enrichment Analysis (GSEA) based on Gene Ontology (GO) (Ashburner et al., 2000; Consortium, 2020) implemented in the goseq R package (Young et al., 2010) and metabolic pathways were defined by Kyoto Encyclopedia of Genes and Genomes (KEGG) (Kanehisa, 2000) implemented in the clusterProfiler R package (Yu et al., 2012).

#### 2.2.3 Omics integration

The final step of the analysis was the integration of microbiota abundance identified in the 16S rRNA data with the gene expression identified in the RNA-seq data. To study the transcriptome-microbiome interaction we applied the weighted co-expression network analysis implemented in the WGCNA R package (Langfelder and Horvath, 2008). The analysis was split into steps comprising: i. creating a correlation matrix using Pearson’s correlation coefficient between all pairs of genes-genera; ii. Creating adjacency matrix (matrix-based representation of a graph) using the formula: *a*_*mn*_ = |*c*_*mn*_|^*β*^, where *a*_*mn*_ is an adjacency between gene/genus *m* and gene/genus *n, c*_*mn*_ is a Pearson’s correlation coefficient, and *β* is a soft-power threshold determined based on the standard scale-free topology network (Chen and Shi, 2004); iii. transformation of the adjacency matrix into the topological overlap matrix (TOM) which is the matrix of the similarity in terms of the commonality of the connected nodes (Yip and Horvath, 2007); iv. the dynamic tree cutting algorithm was used for hierarchical clustering of TOM into modules, as clusters of highly interconnected genes and genera; iin order to obtain co-expressed modules, the parameters of the algorithm were set to minModuleSize = 20 for the gene/genus dendrogram and minimum height = 0.25 to cut the tree, in order to merge similar modules; v. identification of eigengenes for each module that is expressed by the first principal component of the expression matrix, vi. Pearson’s correlation analysis of eigengenes with phenotypes (with t-test for testing the significance of the correlation coefficient), and finally, vii. identification of hub genes/genera – genes/genera that have the highest correlations with other genes/genera contained within each module. Genes contained within significantly associated modules were then subjected to GSEA.

## 3 RESULTS

### 3.1 Microbiome

The analysis of the 16S rRNA gene allowed us to identify 232 unique genera. The genera, with abundance exceeding 10% in all the samples, were *Clostridium, 57N15*, and *Treponema*. A detailed analysis of microbiota was described by Czech et al. (Czech et al., 2022).

### 3.2 mRNA-seq

The analysis of the RNA-seq data identified 2,035 differentially expressed genes for rectal temperature, 1,886 for drooling score, and 1,958 for respiratory score. The expressions of the majority of those genes were down-regulated – the higher value of phenotypes, the lower expression of these genes. This comprised 85% of down-regulated genes for rectal temperature, 78% for drooling score, and 80% for respiratory score. The most highly up-regulated genes were *ENSBTAG00000048590* (for rectal temperature), *ENSBTAG00000054209* (for respiratory score), and *SLC22A1* (for drooling score), while genes with the highest down-regulated expression were *ENSBTAG00000024272* (for rectal temperature), *ENSBTAG00000050067* (for respiratory score), and *ENSBTAG00000051290* (for drooling score). The 1,851 genes significantly associated with HS were common for all three phenotypes (Figure 1). Next, we performed GSEA in which we identified seven KEGG pathways enriched among significantly expressed genes that were shared between all three phenotypes: herpes simplex virus 1 infection (bta05168), viral protein interaction with cytokine and cytokine receptor (bta04061), chemokine signaling pathway (bta04062), cytokine-cytokine receptor interaction (bta04060), PI3K-Akt signaling pathway (bta04151), antifolate resistance (bta01523), and EGFR tyrosine kinase inhibitor resistance (bta01521). Results of GSEA for KEGGs were visualized in Figure 2. On the plot, we can see that Herpes simplex virus 1 infection is characterized with the lowest P-value of 2.73 · 10^−12^ and also demonstrated the highest gene ratio of significantly associated genes that composed this pathway. Significantly enriched GO terms related to biological processes were identified only for respiratory score and were related to cell surface receptor signaling pathway (GO:0007166), cellular response to endogenous stimulus (GO:0071495), G protein-coupled receptor signaling pathway (GO:0007186), and metal ion transport (GO:0030001) (Figure 3).

**Figure 1.**
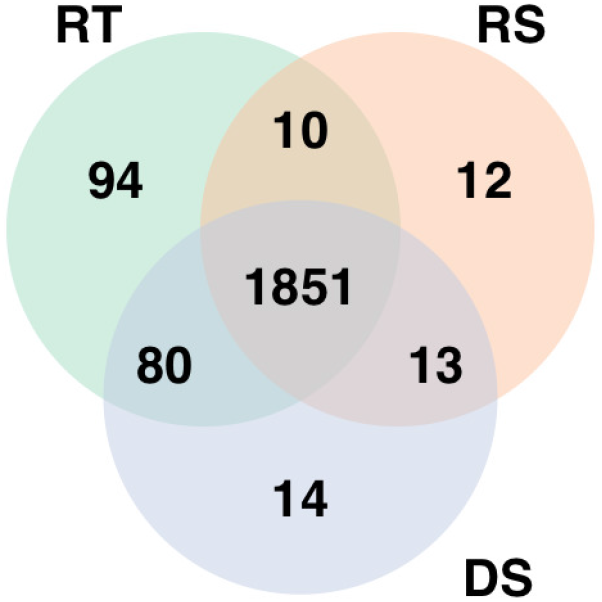
Genes significantly differentially expressed with increasing heat stress for each phenotype. RT denotes rectal temperature, RS respiratory score, and DS drooling score.

**Figure 2.**
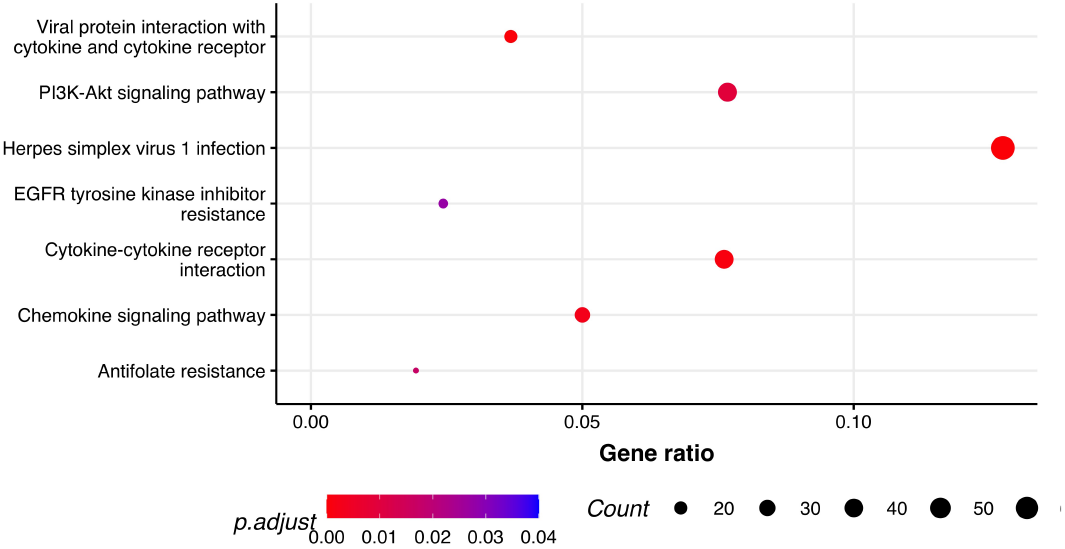
Pathway analysis. Dot plot of the statistically significant KEGG pathways shared between all the phenotypes.

**Figure 3.**
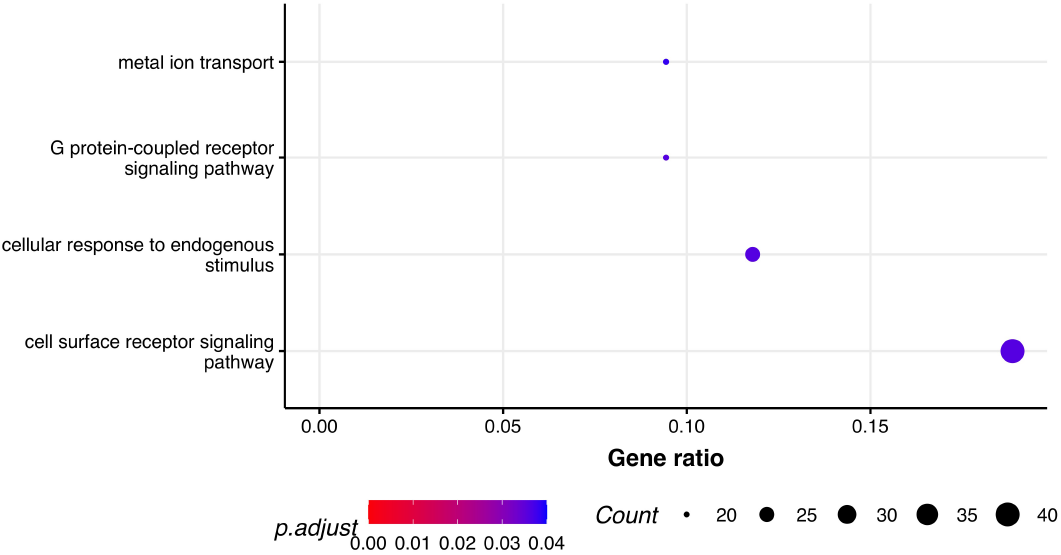
Pathway analysis. Dot plot of the statistically significant GO pathways related to biological processes for respiratory score phenotype.

### 3.3 Omics integration

By applying the steps described in the method section, the weighted co-expression network was generated. The adjacency matrix was created by raising the correlation matrix to the power of 4 (*β* parameter, Figure 4). In the next step, the TOM dissimilarity matrix was computed and used for the hierarchical clustering. Genes and bacteria were clustered into 20 modules, which ranged in size from 36 to 3015 genes/bacteria per module (Fig. 5).

**Figure 4.**
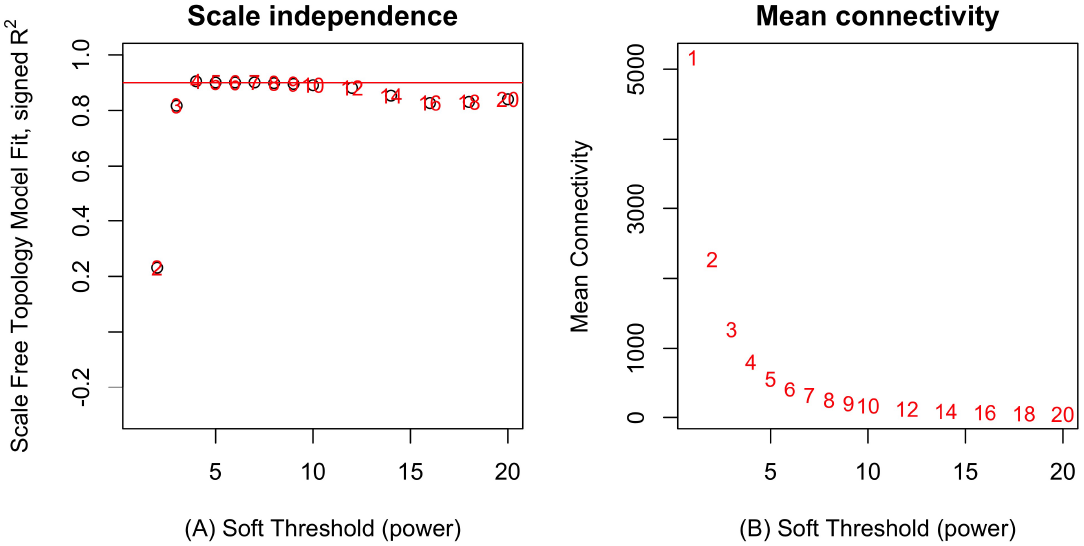
Network topology analysis for soft-thresholding powers in WGCNA – scale-free fit index for different powers (A) and mean connectivity analysis for different soft-thresholding powers (B).

**Figure 5.**
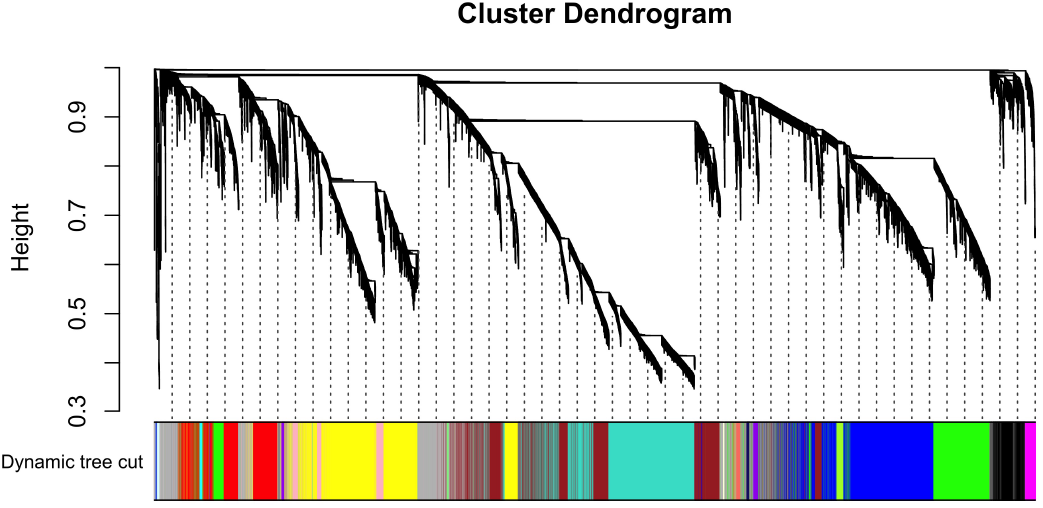
Hierarchical cluster tree of co-expressed genes and bacteria.

The effect of each gene was expressed by the eigengene value, and the correlation of each eigengene with each HS phenotype was calculated (Figure 6). Three modules showed significant correlations with rectal temperature (MEtan, MElightycan, and MEroyalblue). Module MEtan consists of 129 genes but no bacterial genera, MElightycan of 26 genes and 26 bacterial genera, and module MEroyalblue of 2 genes and 34 bacterial genera.

**Figure 6.**
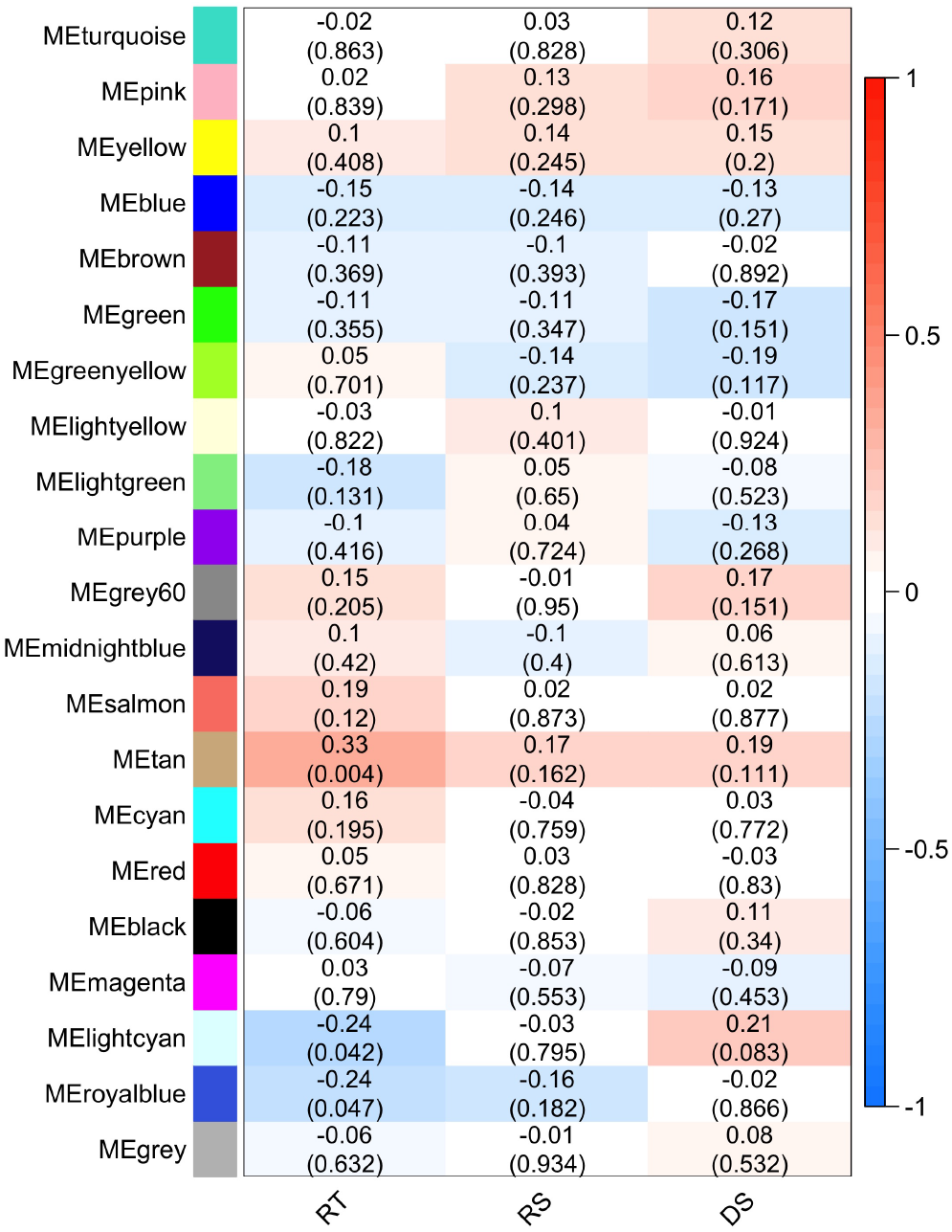
Module-trait associations. Each row corresponds to a module eigengene, column to a trait – rectal temperature (RT), respiratory score (RS), and drooling score (DS). Each cell contains the corresponding Pearson’s correlation and P-value.

Further, we identified hub genes/bacterial genera representative for each of the three modules: *CSF3R* gene in MEtan, *Lactococcus* bacteria in MElightcyan, and *Rhizobium* bacteria in MEroyalblue. There was no overlap between genes contained within the significant modules and in the differential gene expression analysis. We annotated all genes in significant modules to GO terms and KEGG pathways. MEtan module was enriched in a pathway related to *Pertussis* and *Salmonella* infection (bta05133 and bta05132, respectively) and in GO terms described the cellular response to organic substance (GO:0071310), response to oxygen-containing compound (GO:1901700), cellular response to lipid (GO:0071396), and cellular response to lipopolysaccharide (GO:0071222). The other two modules showed no significant enrichment of KEGG and GO terms.

### 3.4 Discussion

This experiment is one of the first investigation in which the combined data of host transcriptome and microbiota were used together to study heat stress in cattle. The direction of changes in cows under heat stress was identified on the level of gene expression alteration as well as on the level of the interaction with microbiota. Heat stress is undoubtedly a complex process that scientists today must face in order to protect animals. However, due to its physiological complexity, we are not able to assess in detail the changes in molecular mechanisms underlying HS response in livestock. The progressive development of molecular biology and bioinformatics allows for a broader look into changes in organisms, allowing simultaneous insight into the cell at virtually every stage of its life cycle. The RNA-seq technology has become a very powerful method for identifying candidate genes associated with complex traits. Already in 2020 Garner et al. (Garner et al., 2020) identified *BDKRB1* and *SNORA19* as potential candidate genes related to HS. Sigdel (Sigdel et al., 2019) reported *HSF1, MAPK8IP1*, and *CDKN1B* as genes responsible for thermotolerance in dairy cattle. Moreover, Diaz (Diaz et al., 2021) identified five genes: *E2F8, GATAD2B, BHLHE41, FBXO44*, and *RAB39B* which were significantly associated with HS. In this study, it was found that the gene *RAB39B* was significantly associated with three phenotypes which included rectal temperature, drooling, and respiratory scores.

For the microbiome, Sales (Sales et al., 2021) identified four bacterial genera related to HS – *Flavonifractor, Treponema, Ruminococcus*, and *Carnobacterium*. In this study, it was identified that HS inhibits gene expression of several genes that might be related to the reduction of energy during overheating. Because HS appears to be a physiologically complex phenomenon, the multiomics approach that accounts not only for alteration in gene expression and changes in the microbiome composition but also for the interaction between them is an important method. In this study, which is follow-up analysis, genes and pathways were identified that are significantly associated with HS phenotypes. Additionally, interactions involving mRNA levels and microbiota in cattle were analyzed. Although the overlap between these findings and the microbiome and genes related to HS reported in the literature is constrained to only *RAB39B*, therefore it is hypothesized that this approach which focused on the interaction between microbiome and host genetics was able to identify new components of the HS response that have been missed in the single omics analyses. A loss of interaction under increased HS was observed. In two out of three significant modules, bacteria played a key role in the regulation of gene expression and controlled the abundance of other bacteria. In particular gene *CSF3R* that was identified as the only hub gene in all significantly associated coexpression modules. Currently the importance of this gene in the context of heat stress in cattle has not been reported yet. However, in human genetics this gene is associated with congenital neutropenia (Triot et al., 2014). Park (Park et al., 2021a) already reported that HS may affect neutrophil phagocytosis. It may indicates that gene *CSF3R* might be strictly associated with both neutrophils and heat stress response in cattle. Other significant hubs were represented by bacteria. *Lactococcus* bacteria that was identified as the hub of MElightcyan module was already indicated in the literature as a genus associated with bovine mastitis (Rodrigues et al., 2016). This observation stressed the important role of the gut microbiome in the regulation of gene expression.

## AUTHOR CONTRIBUTIONS

B.C., Y.W. and J.S conceived and conducted the experiment, B.C. analyzed the results and wrote the manuscript in consultation with J.S., Y.W., K.W., H.L., L.H. All authors reviewed the manuscript. All authors read and approved the final manuscript.

## ACKNOWLEDGMENTS

Calculations have been carried out using resources provided by Wroclaw Centre for Networking and Supercomputing (http://wcss.pl), grant No. 509.

Computations were carried out at the Poznan Supercomputing and Networking Centre.

## DATA AVAILABILITY STATEMENT

The sequence data are available in the NCBI Sequence Read Archive with accession number SRP202074. Other datasets generated and/or analyzed during the current study are not publicly available due to institutional constraints but are available from Yachun Wang on reasonable request.

